# Designing flow regimes to support entire river ecosystems

**DOI:** 10.1101/2020.01.09.901009

**Authors:** Jonathan D. Tonkin, Julian D. Olden, David. M. Merritt, Lindsay V. Reynolds, Jane S. Rogosch, David A. Lytle

## Abstract

Overcoming the challenges of water scarcity will require creative approaches to flow management and modeling approaches that forecast the effects of management actions on multiple ecosystem components simultaneously. Using a mechanistic multispecies modeling approach, we investigated the cross-ecosystem effects of environmental flow regimes designed for specific ecosystem outcomes. We reveal tradeoffs associated with flow regimes targeting riparian vegetation, fishes, and invertebrates. The different frequencies associated with each flow regime in some cases caused non-target ecosystem components to become locally extirpated within short (decadal) timespans. By incorporating multiple flow frequencies (from intraannual-scale pulses to large decadal-scale floods), the natural flow regime enabled a balanced but sub-optimal response of the three ecosystem components (mean 72% of designer flow). Although returning to a natural flow regime may not be possible in highly managed rivers, novel flow regimes must incorporate diverse frequencies inherent to such a regime and accommodate the sometimes conflicting requirements of different taxa at different times.

## Introduction

Dams and other types of human infrastructure have modified river hydrology globally and continue to do so at an unprecedented rate (Grill et al., 2019). Alteration of river flows comes at a major cost for the biota that inhabit freshwater and riparian (hereafter “river”) ecosystems (Tonkin et al., 2018; Bunn and Arthington, 2002), threatening the countless ecosystem services they provide (Auerbach et al., 2014). Maintaining functional river ecosystems under uncertain hydroclimatic futures presents a major management challenge for both existing and planned dam projects (Tonkin et al., 2019; Palmer and Ruhi, 2019; Horne et al., 2019), requiring the consideration of flow prescriptions that target the health of downstream ecosystems (Acreman et al., 2014).

Environmental flows are increasingly used to help minimize the detrimental effects of dam management on river biota (Yarnell et al., 2015; Poff and Matthews, 2013). Designer environmental flows range from single events designed to achieve a specific goal, such as a flood for mobilizing sediment, to entire flow regimes designed to accommodate multiple ecosystem needs (Acreman et al., 2014). Although increasing attention has focused on tradeoffs between ecosystem and multiple human needs (domestic, agriculture, hydropower) in flow designs (e.g. Sabo et al., 2017; Chen and Olden, 2017; Tickner et al., 2017), the large majority of these management decisions are based on a narrow perspective of the ecosystem. In practice, most environmental flow programs target a few important species or a particular component of the river ecosystem, such as recruitment of riparian vegetation or spawning of native fish (Olden et al., 2014) without directly considering secondary effects on other components of the ecosystem. For instance, a flow regime designed to maximize fish abundance or diversity may have unintended less beneficial, or even detrimental, effects for native riparian plants. This presents the question: Does flow management designed to benefit one important component of the river ecosystem simultaneously protect other components, or does it involve ecological tradeoffs that compromise other ecosystem components? In reality, such trade-offs will be hard to avoid, but their magnitude will likely depend on the specific target of the management action.

Calls for holistic ecosystem approaches to support sustainable riverine management continue to mount, yet robust quantitative models to underpin such efforts remain scarce (Poff and Olden, 2017). Here, we examined the responses to designer flows targeting three ubiquitous but distinct components of river ecosystems: riparian vegetation, fishes, and aquatic invertebrates. Using a mechanistic, multispecies modeling approach that links population dynamics and hydrology (WebTable 1), we designed flow regimes to maximize management outcomes for specific targets within each of the three ecosystem components: a riparian tree (cottonwood, *Populus fremontii*), native freshwater fishes, and terrestrially-available aquatic invertebrates. These flow regimes had characteristic flow event frequencies ranging across intraannual- to decadal-scales. The modeling approach permitted us to simultaneously design flow regimes for a single ecosystem component as well as quantify the associated synergies or tradeoffs across other ecosystem components. These approaches enable an assessment of the potential ecological benefits or deficits of designer flow prescriptions for whole river ecosystems.

## Methods

### Management targets

Using species common to the Colorado River Basin of the southwestern United States, we defined management targets that natural resource managers often seek to maximize downstream from large dams. These outcomes relate to three components of river ecosystems: cottonwood tree coverage as a percent of riparian carrying capacity, native fish species biomass as a percent of total carrying capacity (including nonnative fishes), and abundance of terrestrially-available benthic invertebrates (hereafter “aquatic invertebrates”). Cottonwood is an important native riparian tree that suffers from the effects of flow regime modification and competition with nonnatives such as tamarisk (Merritt and Poff, 2010). Native fish species are a focal management target because they face an uncertain future resulting from altered hydrology and impacts from nonnative species (Chen and Olden, 2017). Terrestrially-available aquatic invertebrates serve as aquatic prey for fish and, upon emergence as winged adults, as terrestrial prey for birds, bats, lizards and other riparian animals (Baxter et al., 2005). Using these targets, coupled with mechanistic population models, we identified flow regimes that maximized the value of each ecosystem component and explored tradeoffs in achieving positive outcomes across all three components by projecting the flow time-series for up to 200 years into the future. We ran simulations on a river modeled after a generalized tributary of the Colorado River. We also examined a natural flow regime scenario using a hydrograph derived from a large free-flowing river supporting riparian vegetation, fish and aquatic invertebrate assets (upper Verde River, Arizona, USA) and quantified the community-wide population responses across ecosystem components (Fig. 1). Detailed methods are presented in WebPanel 1.

**Figure 1:**
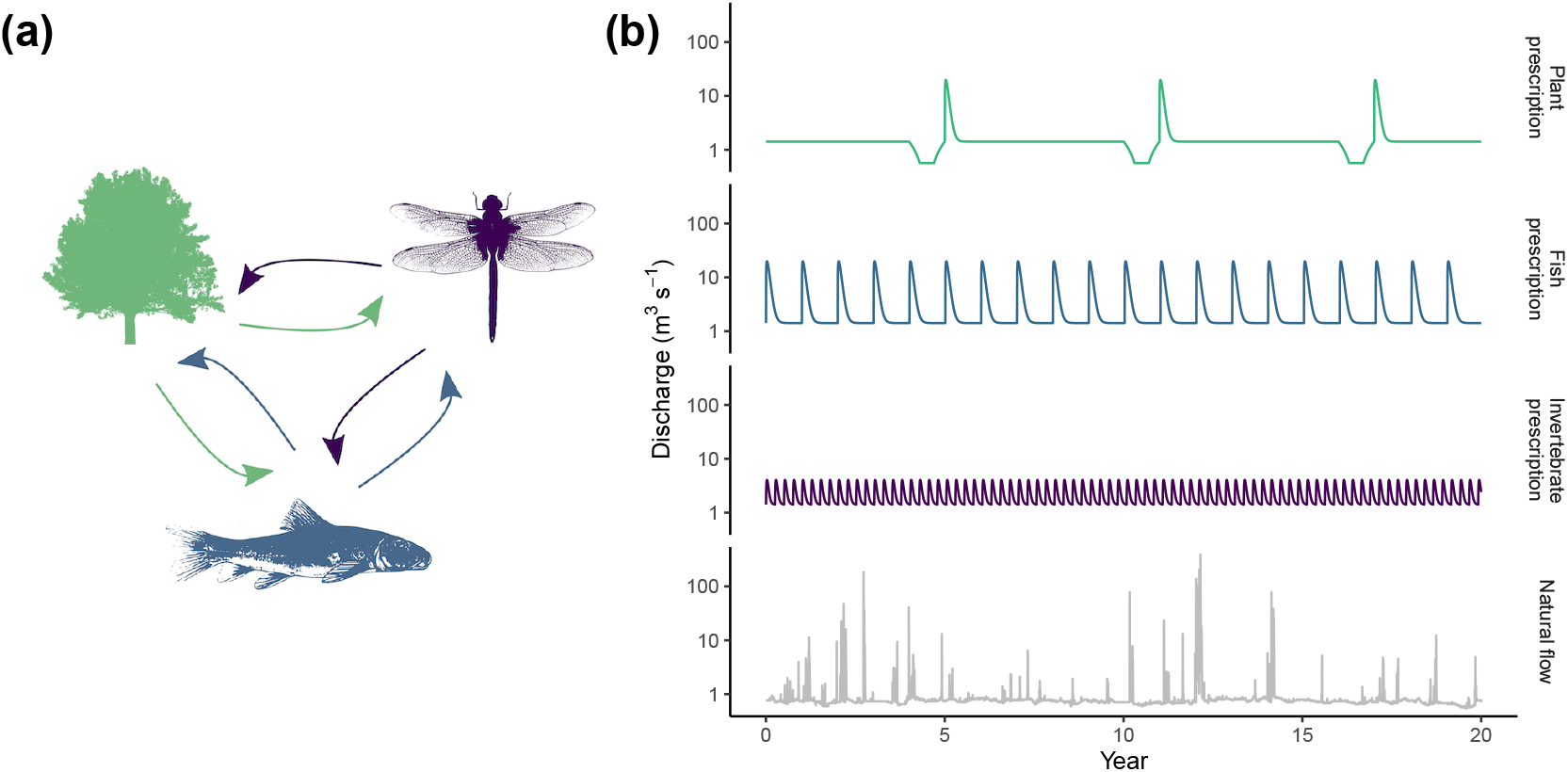
The modeled ecosystem components and flow regimes. A. The three ecosystem components examined: riparian plants (target: % cottonwoods), fish (target: % native species), and benthic invertebrates (target: % terrestrially-available taxa). Arrows represent potential tradeoffs associated with specific flow prescriptions. B. A schematic of the three flow regimes designed for each target ecosystem component. We searched flow parameter space for optimal sequences of flow events that maximized each of the management targets. The plant and fish models are based on year-types and the invertebrate model responds to individual flow events, thus the ‘hydrographs’ are for visual purposes only (e.g. the plant prescription shows a single large flood event every 6 years, preceded by a drought year). The bottom panel represents the historical hydrograph taken from the Verde River, Arizona, USA. Note the logarithmic scale on the y-axis.

### Modeling frameworks

We modeled the three ecosystem components using three independent existing models parameterized with empirical data. Riparian vegetation and fishes were modeled using coupled, stage-structured matrix population models projected at annual time steps (Lytle et al., 2017; Rogosch et al., 2019). Informed by empirical data from a variety of sources, these models link the flow regime directly with population dynamics in a coupled framework that enables an understanding of whole-community dynamics and can incorporate stochasticity by taking random draws from a sequence of river flow year types. Floods and droughts interact with vital rates, affecting population sizes, which opens vacant space (vegetation) or biomass (fish) for recruitment during the next year if conditions are met. Both models have demonstrated a strong ability to recover known patterns on the landscape via tests against empirical data (Lytle et al., 2017; Rogosch et al., 2019).

The plant community comprised five taxa with six stage classes from seedling to reproductive adult: cottonwoods (*Populus deltoides*), tamarisk (*Tamarix ramosissima*), willow (*Salix exigua*), meadow grasses, and sagebrush (*Artemisia tridentata*). These taxa are representative of dominant groups across dryland regions. The fish community comprised seven species, each with three stage classes. Three species are native to the southwest USA: desert sucker (*Catostomus clarki*; CACL), Sonora sucker (*Catostomus insignis*; CAIN) and roundtail chub (*Gila robusta*; GIRO); and four are non-native: yellow bullhead (*Ameiurus natalis*; AMNA), green sunfish (*Lepomis cyanellus*; LECY), smallmouth bass (*Micropterus dolomieu*; MIDO), and red shiner (*Cyprinella lutrensis*; CYLU).

Benthic invertebrates, which experience population dynamics at intra-annual timescales, were modeled using a form of the continuous logistic growth model that enables carrying capacity (*K*) to fluctuate through time (McMullen et al., 2017). Carrying capacity, in this case, responds to flood events. For flood-adapted species, carrying capacity is highest immediately post-flood. The magnitude of a flood pulse determines the magnitude of change in *K*, and this relationship can be modeled for events of any magnitude and for multiple, repeated events. Contrary to the fish and riparian plant models, this model operates on individual populations in that the populations do not share a finite resource such as space or biomass: each population has its own carrying capacity. As representatives of a diverse range of aquatic invertebrate life histories, we modeled three invertebrate taxa: a fast life-cycle, flood-adapted mayfly, *Fallceon* spp. (Ephemeroptera: Baetidae); a slow life-cycle, flood-adapted dragonfly *Progomphus* spp. (Odonata: Gomphidae); and a flood-averse ostracod seed shrimp (Crustacea: Ostracoda) (see WebPanel 1). Additionally, the mayfly and dragonfly are important resources in both aquatic and terrestrial food-webs owing to their aerial adult stages. Thus, our management scenario seeks to maximize the mayfly and stonefly population sizes, and minimize the ostracods.

Vital rates for all species included in the models were obtained from independent sources in the literature and from field studies (see WebPanel 1 for details). Vital rates included stage-specific mortality rates in response to flow events for fish and riparian plants, and population growth rates and flow-specific mortality rates for aquatic invertebrates.

### Natural hydrograph details

Although our approach was simulation-based (with parameterizations from field data) in a generalized river in the southwestern United States, we used a real flow regime to generate a natural hydrograph vector. To do this, we sourced a 45-year (1964-2008) historical hydrograph from the upper Verde River (USGS gauge number 09503700) near Paulden, Arizona. All three groups were modeled from this one central flow regime in our natural flow regime scenario, enabling a comparison of community dynamics across the whole ecosystem. For details of conversion of the hydrograph for each model, see WebPanel 1.

### Flow design

We searched flow parameter space for optimal sequences of flow events that maximized each of the management targets associated with riparian plants, native fishes, and aquatic invertebrates (see WebPanel 1). This approach was based on constrained, systematic optimization for a point optima. Although our flow design approach focused on modifying the frequency of different events or year-types, each model incorporated various other components of a flow regime, including magnitude, timing, rate of change, and duration of flow events (Lytle et al., 2017; Rogosch et al., 2019; McMullen et al., 2017). For riparian plants and fishes, this search for an optimal flow design followed a series of steps that incrementally adjusted the frequency of particular year-types. The resulting prescribed flow regime for riparian vegetation consisted of a spring flood every six years, preceded by a drought year, with a series of non-event years in between (WebTable 1). These floods occurred within the spring window that enabled cottonwood and tamarisk to recruit (synchronized with seed release). The resulting prescribed flow regime for native fish percentage consisted of a spring flood every year. In contrast to the plant and fish models, the invertebrate model operates in continuous time. Because the aquatic invertebrates we modeled do not compete directly with each other for a resource such as space or food, we modelled the the three taxa independently so that each species had its own carrying capacity, which we scaled to 100. We sought to maximize the average value of the two terrestrially-available target taxa (i.e. % of *K*) over the 20-year evaluation period. The resulting scenario that maximized the target was four small pulses per year, below the threshold of a flood in either the fish or riparian models. Therefore, the invertebrate prescription results in 100% of years being non-event years for the non-target taxa.

## Results and discussion

We identified specific flow regimes that were highly beneficial to populations of each targeted ecosystem component—cottonwoods, native fishes, and aquatic invertebrates—by maximizing their average population sizes through time (Fig. 1 - 2, WebFigure 1 - 3). These designer flow regimes, optimized separately for each ecosystem target, always outperformed the historical natural regime for the intended ecosystem component, suggesting that artificially-imposed flow regimes can in some instances generate greater population sizes than the natural flow regime. This finding is consistent with modeling studies targeting native fish abundance in the San Juan River, United States, fisheries yield in the Mekong River Basin, and cottonwood population dynamics on the Yampa River, United States (Sabo et al., 2017; Chen and Olden, 2017; Lytle et al., 2017).

**Figure 2:**
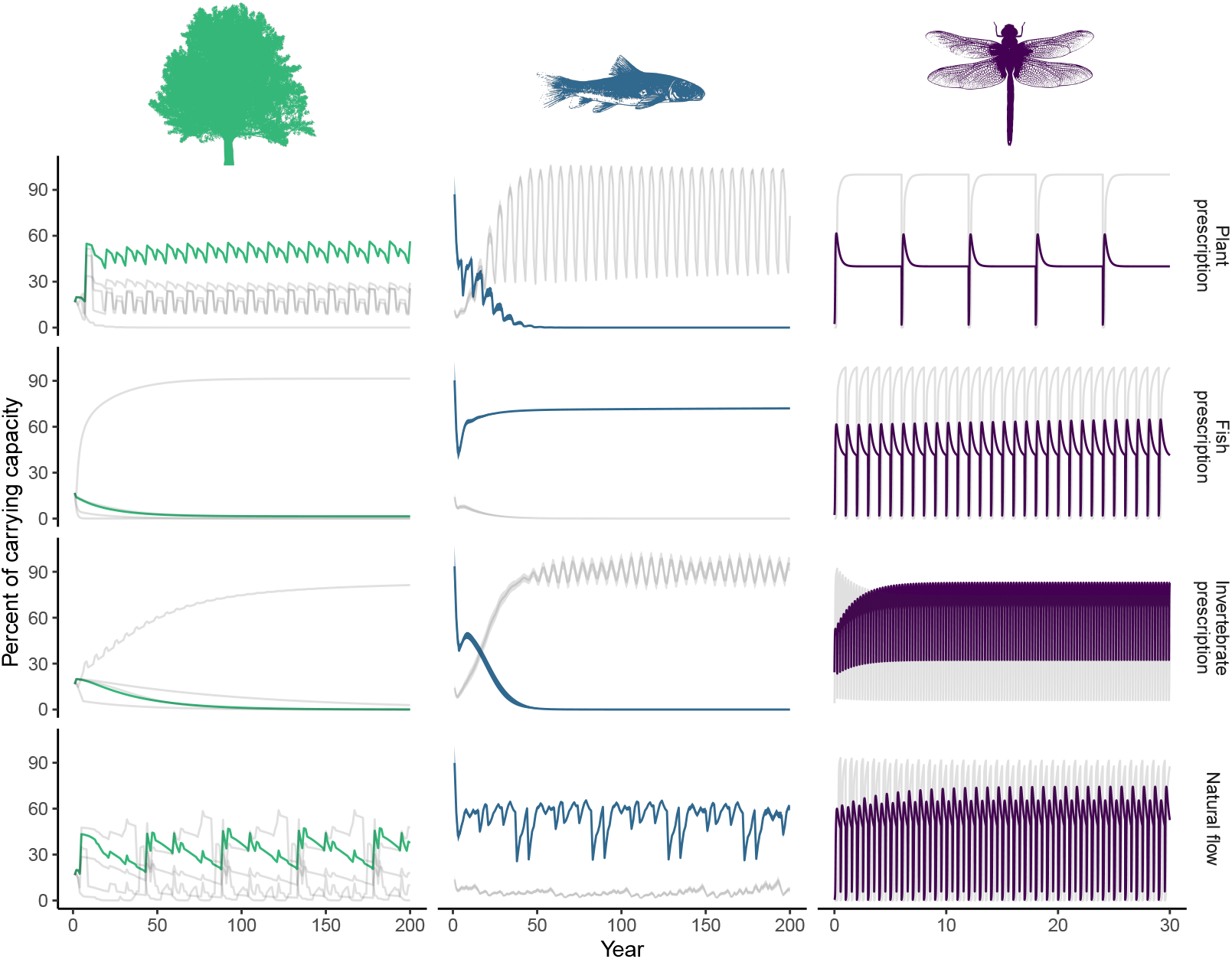
Results of simulations for target and non-target taxa, both within and among ecosystem components. Target taxa (plants: % cottonwood; fish: % native species; invertebrates: % terrestrially-available taxa) are shown as the colored lines and non-target taxa as grey. For plants, the four grey lines represent the four non-target plant taxa individually. For fish, the single grey line represents the four non-native fish species combined (the band around the line represents 2 standard errors around the mean of 100 iterations). For invertebrates, the grey line represents the non-target taxa (ostracods). See Supporting Information for full results. Model evaluation (e.g. Fig. 3) discarded the first ten years as a burn-in period.

**Figure 3:**
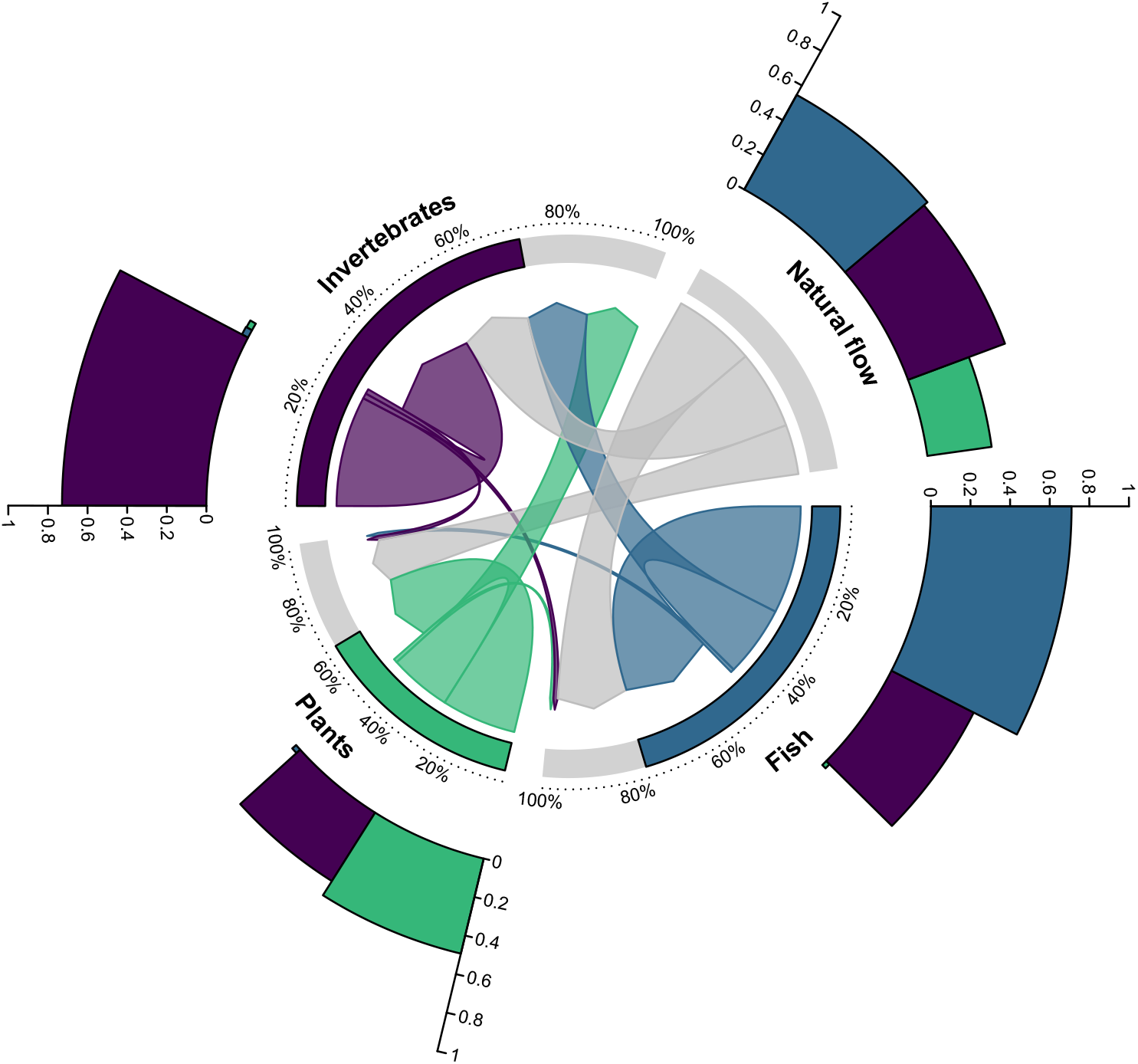
The ecosystem-wide effects of designer and natural flow regimes. Each of the four figure sectors represents a particular flow regime and/or ecosystem component: the natural flow regime and three designer flow regimes (riparian vegetation, fishes, and invertebrates). The effects of designer flow regimes on each component are shown, both targeted (arrow returns to same figure sector; e.g. fish to fish) and non-targeted (arrow from one sector to another; e.g. fish to plants), as well as the natural flow regime. Arrow widths correspond to the relative effects of a particular flow scenario (source of arrow) on all other components (arrow endpoint; larger equal more positive responses). These values are shown as proportions of maximum in the outermost bars (1 = maximum; e.g. native fish biomass 100% of carrying capacity). For instance, when designing flows to most benefit native fishes, fish respond strongly (large arrow and outer bar) but plants perform poorly (small arrow and bar). The bar tracking the circumference of the center arrows represents the difference between the natural and prescribed flow for the group in that sector (given in percentage). For example, under the natural flow regime, fish achieve 77% of the biomass achieved under the designer flow.

However, our results highlight that a narrow management focus on a single taxonomic group, as is commonly the case for environmental flow efforts, may come at a cost for other components of the ecosystem (Fig. 3, WebFigure 4). Each scenario had at least one major losing ecosystem component: cottonwoods declined in the fish-prescribed flows (7% of natural flow %*K*), native fish abundance declined in the vegetation-prescribed flows (5% of natural flow %*K*), and both cottonwoods (10% of natural flow %*K*) and native fishes (6% of natural flow %*K*) declined in the invertebrate-prescribed flows. Thus, major ecological deficits appear to accompany environmental flow regimes that target a single ecosystem outcome.

Each designer flow regime had a characteristic temporal frequency, reflecting the varying biology of the three ecosystem components: approximately decadal or half-decadal timescale flow events for riparian vegetation, annual or near-annual for fishes, and intra-annual for invertebrates (Fig. 1 - 2, WebTable 1). Cottonwood thrive under regimes with large recruitment floods approximately every six years followed by growth years (i.e. non-event years), with a drought year included to limit population growth of drought-intolerant competitor species (Lytle et al., 2017). Native fishes prosper under flow regimes with more frequent and reliable (i.e. annual) spawning floods and no drought (Rogosch et al., 2019). Thus, bigger floods are beneficial for vegetation and fishes, but the timescale of response differs. By contrast, the pulses required for maintaining aquatic invertebrates are insufficiently large to exert a positive benefit for native fishes or cottonwood; this taxonomic group responds best to flow regimes comprising many regular small pulses to maintain shallow riffle habitat. Without regular small pulses (larger pulses may also be incorporated), slower-water specialist invertebrates, many of which do not have a terrestrial lifecycle phase, become dominant (McMullen et al., 2017). In summary, various temporal frequencies are thus fundamental aspects of a flow regime designed for the benefit of an entire ecosystem—a characteristic of the historical natural flow regime that is essential for the vitality of rivers (Fig. 1) (Poff et al., 1997; Naiman et al., 2008; Tonkin et al., 2019). Administering such ecosystem-level designer flows may be challenging if single sensitive (threatened, endangered, or red listed) species provide the impetus (legal mandate) for implementation of designer flows.

Contrary to the designer flow regimes, the natural flow regime scenario resulted in species persistence for all ecosystem components, although population sizes were never as large as those achievable under designer flow regimes (vegetation: 66% of designer flow; fishes: 77%; invertebrates: 72%) (Fig. 2). We attribute this to the fact that most natural flow regimes exhibit an array of hydrologic events at multiple temporal frequencies (from intraannual to inter-decadal), thereby satisfying the ecological needs of diverse biological groups with sometimes conflicting requirements (Fig. 1 - 2). Most dam operations fail to provide this diverse portfolio of flows that are important for recruitment, migration, spawning and juvenile rearing across a broad array of taxa (Palmer and Ruhi, 2019). Organisms have evolved life histories to capitalize on natural cycles of flooding and drought (Lytle and Poff, 2004). However, the evolutionary fine-tuning, and potential for rapid evolution, of entire ecosystems to the natural flow regime remains an important topic of inquiry, as does the interaction among the different ecosystem components that we have yet to consider (e.g. invertebrates are a food source for fish, plants provide organic matter for invertebrates). Maintaining these cycles is fundamental to the maintenance of diverse and resilient communities into the future (Tonkin et al., 2018). Flooding also plays a critical role in maintaining functional river geomorphology by creating and maintaining critical off-channel habitats, and mobilizing sediment, woody debris and essential nutrients (Yarnell et al., 2015). Whether maintaining such variability is possible with environmental flows remains to be seen given the rapidly shifting state of river flows worldwide (Poff, 2018; Poff and Olden, 2017; Tonkin et al., 2019).

Interannual variability in flows that are supported by natural flow regimes enables the persistence of multiple species across diverse taxonomies (Fig. 2). Simply put, some flows benefit particular suites of species in certain years to the detriment of others, but gains made during these periods enable their persistence and coexistence with other species, often through unfavorable periods, over long time-scales (Tonkin et al., 2017; Ruhí et al., 2016). Through time, the full dynamism of river hydrology accommodates higher temporal diversity, emphasizing the importance of taking a functional whole-regime approach to designing and prescribing environmental flows (Yarnell et al., 2015). In long-lived species, such as riparian vegetation, desired outcomes may only manifest in response to flow prescriptions that operate over multiple years or decades. Bottomline: plants and animals are committed to long-term flow regimes; humans similarly need to be committed to long-term flow management.

The mechanisms that produced undesired non-target outcomes were specific to each ecosystem component. For fishes, native species were projected to become locally extirpated within approximately 50 years due to a complete lack of flood recruitment events reflected in the invertebrate flow prescription, or a combination of droughts with too-infrequent flood events reflected in the riparian flow prescription, both of which allowed non-native fishes to dominate the community (Fig. 2, WebFigure 2). This finding is supported by empirical research in the American Southwest (Chen and Olden, 2017; Ruhí et al., 2016; Rogosch et al., 2019). Cottonwoods collapsed in response to native fish-prescribed flows due to phreatophytic, flood-tolerant willow species dominating at high flood frequencies. In response to a lack of recruitment flood events under the invertebrate flow prescription, cottonwoods were replaced by non-riparian upland species such as sagebrush (Fig. 2, WebFigure 1), representing a loss of riparian trees and shrubs that comprise essential and high quality habitat for diverse terrestrial fauna (Merritt and Bateman, 2012). The target invertebrates exhibited much greater fluctuations in population abundance than fish or vegetation (Fig. 2, WebFigure 3). Both the fish and riparian prescriptions did not meet the needs of the aquatic invertebrates due to a lack of regularly-spaced pulses required to maintain open habitat for the two target taxa: a flood-resilient mayfly *Fallceon* spp. (Ephemeroptera: Baetidae), and a flood-resistant dragonfly *Progomphus* spp. (Odonata: Gomphidae) (McMullen et al., 2017). These clear tradeoffs reflect situations where flows are targeted not for an entire community response (e.g. community evenness), but a specific component of each community. Different outcomes may be apparent with alternative ecological targets, but the important implication of this research is that narrowly targeting individual ecological outcomes with specific flows may have broader ecosystem-wide negative impacts.

Different frequencies of response to flows among taxa presents unique challenges when setting out to optimally manage rivers in an environmental flows context. Experimental flood programs have in some cases demonstrated benefits to non-target ecosystem components (e.g. River Spöl, Switzerland: Robinson et al. (2018); Bill Williams River, USA: Shafroth et al. (2010)). However, the potential ecosystem-wide impacts of narrowly prescribed flow regimes emphasizes the need to consider entire ecosystems as the ultimate management goal, rather than focus on single physical (e.g. sediment) or biological (e.g. fish) outcomes (Olden et al., 2014). Considering whole-ecosystem integrity will inevitably require mimicking some functional components of natural hydrologic variability (Yarnell et al., 2015), most notably the presence of multiple frequencies and magnitudes of flow events over extended timescales, while minimizing unnatural frequencies such as daily hydropeaking (Kennedy et al., 2016). However, hydroclimatic nonstationarity, where the envelope of variability in which a river flow regime fluctuates no longer remains fixed (Milly et al., 2008), means returning to the inherently dynamic natural flow regime as a management target may no longer be the most beneficial option (Poff, 2018; Acreman et al., 2014; Tonkin et al., 2019). Overcoming the physical limits of dam operations under nonstationarity, therefore, requires creative approaches to flow management (Poff and Olden, 2017) and a coherent modeling approach that forecasts the effects of management actions on multiple ecosystem components simultaneously.

The unprecedented magnitude at which river flows are being altered across the developing world, combined with the already large proportion of dammed rivers in the developed world, puts into question the long-term sustainability of freshwater ecosystems (Poff and Matthews, 2013; Grill et al., 2019). Environmental flow regimes targeting single ecological outcomes may help to alleviate some of these detrimental effects, but we urge caution in their application due to the potential of unintended collateral impacts on other components of the ecosystem. How can entire ecosystems be better considered in modern day flow management strategies? We assert that designing flows for the benefit of entire ecosystems requires long-term perspectives that embrace hydrologic dynamism involving critical flow events that occur at multiple temporal frequencies. Modeling tools, whether mechanistic or statistical, are critical for better managing dammed rivers, particularly when embedded in an iterative cycle that includes prediction, testing and improvement as new evidence arrives (Tonkin et al., 2019; Dietze et al., 2018; Konrad et al., 2011). Although returning to the historical natural flow regime in managed rivers is an increasingly distant option in a nonstationary world (Tonkin et al., 2019), environmental flows must remain founded on the principles of the natural flow regime paradigm by incorporating the variability to which native species and communities have evolved.

## Acknowledgements

We thank two anonymous reviewers for helping improve the manuscript. Funding was provided in part by the US Department of Defense (SERDP RC-2511) and the US Department of Agriculture Forest Service. JDT is supported by a Rutherford Discovery Fellowship administered by the Royal Society Te Apaārangi (RDF-18-UOC-007).

## Code and data availability

Code and data used in the analysis are available at https://github.com/jdtonkin/flow-tradeoffs.

