## Supporting information for "Designing flow regimes to support entire river ecosystems"

### JD Tonkin *et al.* – Supporting Information

### WebPanel 1. Methods

#### Target scenarios

Using species common to rivers in the Colorado Basin of the southwestern United States, we defined biodiversity target results that managers may seek to maximize downstream from a large dam for three components of a river-riparian (hereafter "river") ecosystem: cottonwood tree coverage as a percent of reach-wide riparian community carrying capacity, native fish species biomass as a percent of reach-wide fish community carrying capacity, and terrestrially-available benthic invertebrates as a percent of population carrying capacity. Using these targets, coupled with mechanistic population models, we designed flow regimes that maximized the value of the target. We then explored the population- and community-level impacts of these flow regimes on other components of the ecosystem by projecting each flow regime up to 200 years into the future and quantifying the outcome. We ran simulations on a river modeled after a generalized tributary of the Colorado River. We also examined a natural flow regime scenario using a hydrograph derived from a large free-flowing river in the southwestern USA (Verde River, Arizona) and quantified the relative differences and tradeoffs across the ecosystem components.

#### Modeling frameworks

We modeled the three ecosystem components using three independent existing models (Lytle et al., 2017; McMullen et al., 2017; Rogosch et al., 2019). Riparian vegetation and fishes were modeled using an application of coupled, stage-structured matrix population models (Lytle et al., 2017; Rogosch et al., 2019). These models link the flow regime directly with population dynamics in a coupled framework that enables an understanding of whole-community dynamics and can incorporate stochasticity by taking random draws from a sequence of river flow year types. The models operate at the reach scale and share a central dependency assumption that represents the whole-community carrying capacity for which the organisms exploitatively compete. A unique attribute of these models is their ability to identify how complex species interactions change under novel environmental conditions, which allowed us to quantify species performance under all possible environmental states (Tonkin et al., 2018). Space (areal extent) in the riparian landscape is the finite element for riparian plants (Lytle et al., 2017), whereas maximum total biomass for each reach is the finite element for fishes (Rogosch et al., 2019). The models employ different methods to realize density dependence, which can be found in the relevant publications (Lytle et al., 2017; Rogosch et al., 2019;

McMullen et al., 2017). Floods and droughts interact with vital rates, affecting population sizes, which opens vacant space or biomass for recruitment during the next year if conditions are met. Both the vegetation and fish models have demonstrated a strong ability to recover observed trends via tests against empirical data (Lytle et al., 2017; Rogosch et al., 2019).

The plant community comprised five taxa with six stage classes from seedling to reproductive adult: cottonwoods (*Populus deltoides*), tamarisk (*Tamarix ramosissima*), willow (*Salix exigua*), meadow grasses, and sagebrush (*Artemisia tridentata*). These taxa are representative of dominant groups across dryland regions. Each taxon $j$ is described by a stage-based matrix $\boldsymbol{n}_j(t+1) = \boldsymbol{A}_j(t)\boldsymbol{n}_j(t)$, where $\boldsymbol{n}_j(t)$ is a vector containing stage abundances and $\boldsymbol{A}_j(t)$ is a set of transition matrices that fluctuate according to variation in the hydrograph (Caswell, 2001). Thus, the population dynamics of each taxa are determined primarily by how their vital rates (stage-specific flood and drought mortality, fecundity, self-thinning) are affected by annual cycles of flooding and drought. Vital rates were sourced from a variety of locations including aerial photograph time-series, literature, experimentation (e.g. Cooper et al., 1999) and field surveys. Detailed definitions of the vital rate parameters and model structure can be found in Lytle et al. (2017). The plant model is fully ergodic, so we ran a single simulation for all scenarios.

The fish community comprised seven species, each with three stage classes, covering a range of body sizes and representing the major life-history tradeoffs among growth, survival, and reproduction (Olden et al., 2006). The community comprised three species native to the southwest USA: desert sucker (*Catostomus clarki*; CACL), Sonora sucker (*Catostomus insignis*; CAIN) and roundtail chub (*Gila robusta*; GIRO); and four non-native species: yellow bullhead (*Ameiurus natalis*; AMNA), green sunfish (*Lepomis cyanellus*; LECY), smallmouth bass (*Micropterus dolomieu*; MIDO), and red shiner (*Cyprinella lutrensis*; CYLU). The model conforms to the same general structure as the plant model, with some minor differences in how the vital rates interact with the hydrograph, details of which can be found in Rogosch et al. (2019). Vital rates, including the addition of the Gonadal-Somatic Index to estimate the proportion of egg biomass produced, were sourced from literature reviews. We used 100 iterations for each model run, starting by sampling initial abundance for each species from a negative binomial distribution populated with means and variances from a long-term empirical dataset (seven sites in the upper Verde River, 1994-2008) (Rogosch et al., 2019).

Benthic invertebrates were modeled using a form of the continuous-time logistic growth model that enables carrying capacity ($K$) to fluctuate through time (McMullen et al., 2017). Carrying capacity, in this case, responds to flood events. For flood-adapted species, carrying capacity is highest immediately post-flood. The magnitude of a flood pulse determines the magnitude of change in $K$, and this relationship can be modeled for events of any magnitude and for multiple, repeated events. For flood-averse species, carrying capacity is reduced post-flood and increases with time since flood event. Contrary to the fish and riparian plant models, this model operates on individual populations in that the populations do not share a finite resource such as space or biomass: each population has its own carrying capacity. The details of the model can be found in McMullen et al. (2017).

As representatives of a diverse range of aquatic invertebrate life histories, we modeled three invertebrate taxa: a fast life-cycle, flood-adapted mayfly, *Fallceon* spp. (Ephemeroptera: Baetidae); a slow life-cycle, flood-adapted dragonfly *Progomphus* spp. (Odonata: Gomphidae); and a flood-averse ostracod seed shrimp (Crustacea: Ostracoda). Additionally, the mayfly and dragonfly are important resources in both aquatic and terrestrial food-webs owing to their aerial adult stages. Thus, our management scenario seeks to maximize the mayfly and stonefly population sizes, and minimize the ostracods.

We used the same parameters for each taxon as those in McMullen et al. (2017). Briefly, $K$ for both mayflies and dragonflies was maximized (100) immediately following floods, and declined to $K$ = 40 (40% of maximum possible) after a period of time (depending on the shape parameter $g$, which regulates the rate that $K$ returns to pre-disturbance levels). Both taxa favor open riffle and sandy habitats created by disturbance, which is reduced by encroaching vegetation through time in these dryland rivers. By contrast, $K$ for the ostracod reduced to 40 post-disturbance and increased to 100 after a period of baseflow. Floods reduce preferred habitats of ostracods—slow-water habitats, often created by beaver dam impoundments or large amounts of edge vegetation. The mayfly, dragonfly and ostracod differed in their intrinsic rate of population growth $r$ (mayfly: 0.23; dragonfly: 0.08; ostracod: 0.16) and $h$ (strength of disturbance mortality relationship; mayfly: 0.02; dragonfly: 0.01; ostracod: 0.05). We used days as the unit of time to examine invertebrates. Detailed descriptions can be found in McMullen et al. (2017).

For the sake of simplicity in the invertebrate model, we did not consider in our simulations that 1. there are likely seasonal effects in population dynamics, meaning population growth may be slower during different parts of the year; and 2. with only small floods in our designer flow scenario, other factors such as beaver dam construction, dominance of filamentous green algae, and bed armoring would likely begin to play major roles over time. In this sense, the flows we designed for invertebrates may not be optimal at longer time scales. The major flows that benefit fishes and cottonwoods could be needed for the invertebrates as well to reset the system and reorganize channel and floodplain morphology (Shafroth et al., 2010). Moreover, our modeling does not incorporate negative feedbacks (e.g. low secondary production leading to reduced fish production, despite flow that is tailored towards fishes) that may be present in real-world ecosystems (Anderson et al., 2006). Connecting the three ecosystem components in this way remains an important topic for exploration.

### Hydrograph details

Although our approach was simulation-based in a generalized river in the southwestern United States, we used a real flow regime to generate a natural hydrograph vector. To do this, we sourced a 45-year (1964-2008) historical hydrograph from the upper Verde River (U.S. Geographical Survey gauge number 09503700) near Paulden, Arizona. All three groups were modeled from this one central flow regime, enabling a comparison of community dynamics across the whole ecosystem.

We converted this hydrograph into the format required to inform each of the models. For the plant and fish models, this required converting the continuous time hydrograph into a vector of 'year-types'; the models operate on an annual time-step. These year-types were defined using discharge thresholds for floods and droughts (so the variables are binary). We evaluated the plant and fish simulations by projecting forward 200 years, discarding the first 10 years as burn-in and averaging the remainder of years (see WebFigure 1 and 2). Thus, we first extended the 45-year natural hydrograph by repeating the sequence out to 200 years.

The fish model converts the hydrograph into one of six flow year types: high spring flood, medium spring flood, summer flood, spring/summer flood, drought, and non-event. Having both high spring flood and summer flood occur in one year was a possibility in this model, but all other year types were mutually exclusive. High spring flood events are those that overtop the banks (greater than 19.8 $m^3s^{-1}$, a 4-year return interval), whereas medium spring floods correspond to bankfull events (6.2 $m^3s^{-1}$, 2.5-year return interval). Monsoon season summer floods are roughly equivalent in magnitude to medium spring-flood events. Drought years represent years without floods and with sustained (at least 40 days) low-flow conditions. Non-event years are those that do not meet any of these criteria.

The plant model operates with three independent year types: flood, drought and non-event years. Flood years correspond to the spring high flood years in the fish model, thus representing large bank over-topping floods during the spring recruitment window for cottonwood and tamarisk; "recruitment box model" (Mahoney and Rood, 1998). Likewise, drought years correspond to drought years in the fish model. Non-event years here represent years that are not flood or drought years, thus differing from non-event years in the fish model, since the plant model does not consider medium spring floods or summer floods.

Finally, because the invertebrate model is a continuous-time model that incorporates multiple flood events within a single year, generating a natural flow vector required a different approach than the other models. Broadly, we placed natural flow events into seasonal categories, where the peaks typically occur in these arid southwest USA rivers, to achieve compatibility with the riparian and fish models. First, we took the full historical hydrograph from the Verde River and calculated the mean maximum values for each month across the full sequence to identify seasonal flow peaks. We then identified the seasonal window in which these peaks occurred as they vary annually: spring (larger peak) and late summer/fall monsoon season (smaller peak). We then used this window to search for each of the two peaks, taking the mean maximum flow found in these windows across the full 45-year hydrograph. These two flood pulses represent a single year of the natural flow for the invertebrate model. We repeated this same single year for 30 years to run as the natural flow regime for the invertebrates. We evaluated the invertebrates by projecting forward 30 years, discarding the first 10 years as 'burn-in', and averaging the final 20 years (See WebFigure 3).

#### Flow design

We searched a large amount of alternative parameter space to find the optimal sequence of flow events for each target. For riparian plants and fishes, flow design followed a series of steps that incrementally and systematically adjusted the frequency of particular year-types. For riparian plants, we adjusted the frequency of flood, drought and non-event years until our target (cottonwood percent of $K$) was maximized. This step required us to evaluate community dynamics under a range of flow regimes (sequences and combinations of year-types). This systematic optimization involved first finding the optimum frequency of flood vs. non-event years for cottonwood as we know their life history requires spring floods for recruitment. Using this frequency, we then trialed different additions of drought years until the optimum value for cottonwood was reached. The resulting prescribed flow regime consisted of a spring flood every six years, preceded by a drought year, with a series of non-event years in between (WebTable 1). These floods occurred within the spring window that enabled cottonwood and tamarisk to recruit. Similar results were produced for cottonwoods at frequencies from 2 to 6-year flood return interval, but 6-year return were the optimum frequency. Adding droughts decreases willow, and these drought years were most effective immediately preceding flood years; post-flood drought increased mortality of newly-recruited cottonwood seedlings as drought mortality decreases with life stage.

For fishes, we adjusted the frequency of spring and summer floods, droughts and non-event years. The fish model also has the option for a medium flood in the spring window, but we only focused on large bank-topping floods, which are consistent with the riparian model's spring floods. The resulting prescribed flow regime for native fish percentage consisted of a spring flood every year. Similar to riparian vegetation, multiple settings produced similar results, but annual floods were optimal. The full set of candidate fish flow scenarios consisted of the following: 1. 25% spring flood, 75% non-event; 2. 33% spring flood, 66% non-event; 3. 50% spring flood, 50% non-event; 4. 100% spring flood; 5. 100% spring and summer flood; 6. 50% spring and summer flood, 50% non-event; 7. 50% spring flood, 50% summer flood; 8. 33% spring and summer flood, 66% non-event; 9. 66% spring flood, 33% summer flood; 10. 50% spring and summer flood, 50% spring flood only; 11. 10% spring flood; 12. 20% spring flood,

20% non-event, 60% drought.

In contrast to the plant and fish models, the invertebrate model operates in real time and the three taxa do not interact. Each species has its own carrying capacity, which we scaled to 100. Our target was maximizing the terrestrially available taxa: a dragonfly and a mayfly. To do this, we sought to maximize the average value of these two (i.e. % of $K$) over the 20-year evaluation period. To ensure realism in the allocation of water for the invertebrate prescription, we constrained the maximum number of pulses per year to be four. We sequentially ran through scenarios with a frequency of flood pulses ranging from four per year to one every two years (return interval: 91, 122, 183, 365, and 730 days) and in magnitude from 2.4 to 80.9 $m^3s^{-1}$ (magnitude: 2.4, 3.2, 4.0, 8.1, 16.2, 24.3, 32.4, 40.5, and 80.9 $m^3s^{-1}$). The resulting scenario that maximized the target was four small pulses (4 $m^3s^{-1}$) per year, below the threshold of a flood in either the fish or riparian models. Therefore, invertebrate prescription results in 100% of years being non-event years for the fish and plant targets. The plant- and fish-prescribed models were converted into $m^3s^{-1}$ for the invertebrate model, based on the threshold used to set floods. This resulted in one large flood (19.8 $m^3s^{-1}$) every 2,190 days for the plant prescription and one every 365 days for the fish prescription.

#### Evaluation of model simulations

All model runs were evaluated by discarding the initial ten-year projection as burn-in and averaging the remaining years in each projection: final 20 years of a 30-year projection for invertebrates, final 190 years of a 200-year projection for fish and plants. These values were used to quantify the tradeoff values that went into Fig. 3. All analyses were performed in R 3.3.3 (R Core Team, 2017).

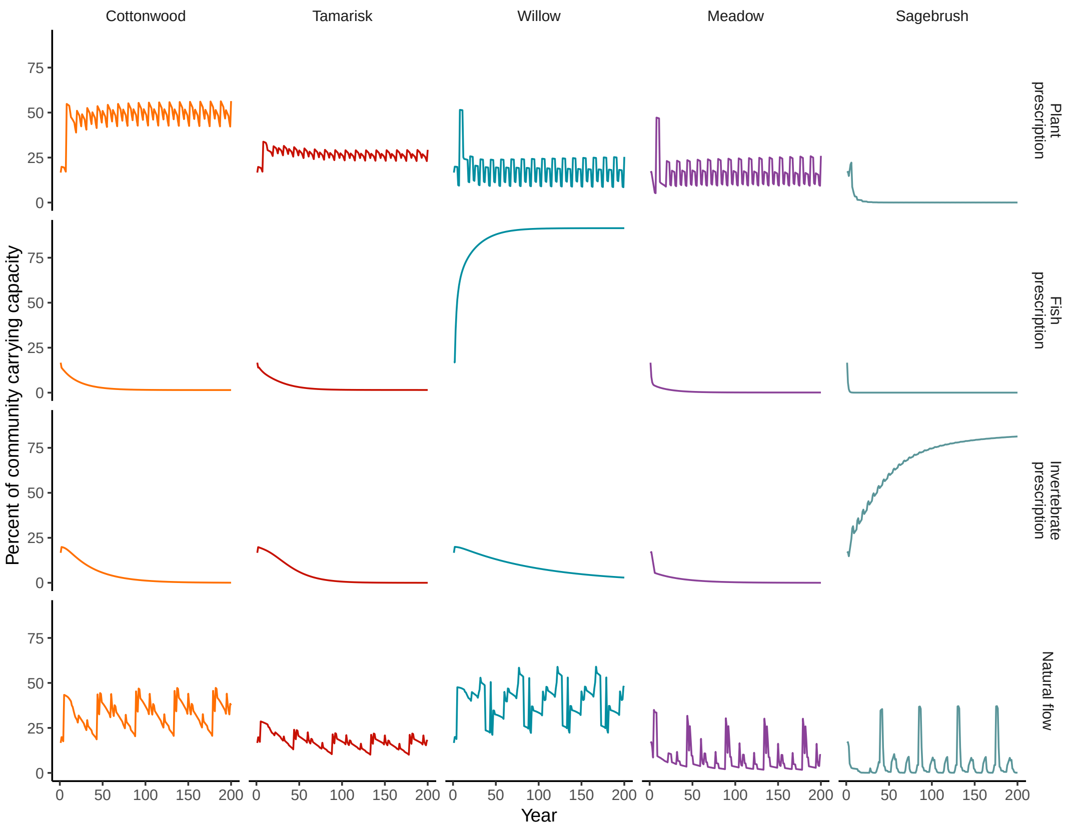


**WebFigure 1:** Results of simulations for the five riparian plant taxa and four flow scenarios. Maximum cottonwood percentage was the flow target.

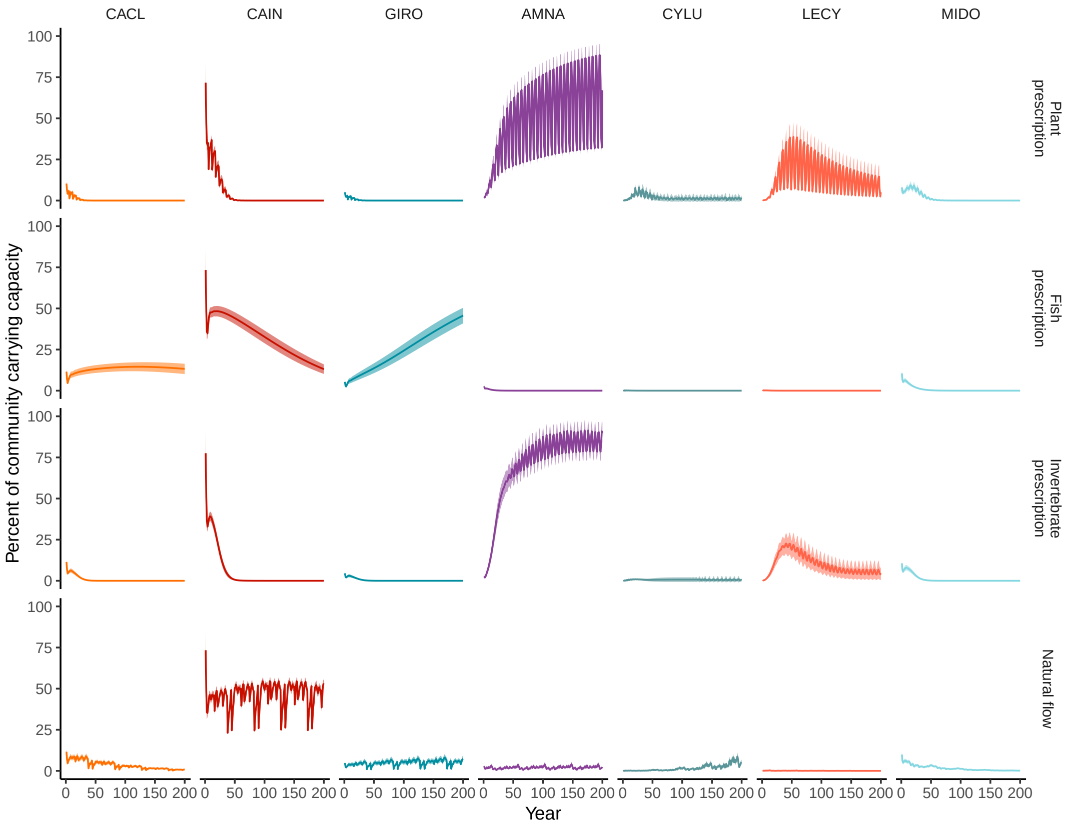


**WebFigure 2:** Results of simulations for the seven fish species and four flow scenarios. The flow target was maximum percent of the three native species to the southwest USA combined: desert sucker (*Catostomus clarki*; CACL), Sonora sucker (*Catostomus insignis*; CAIN) and roundtail chub (*Gila robusta*; GIRO). The four non-native species were: yellow bullhead (*Ameiurus natalis*; AMNA), green sunfish (*Lepomis cyanellus*; LECY), smallmouth bass (*Micropterus dolomieu*; MIDO), and red shiner (*Cyprinella lutrensis*; CYLU). The line represents the mean of 100 iterations and the band around the line represents 2 standard errors.

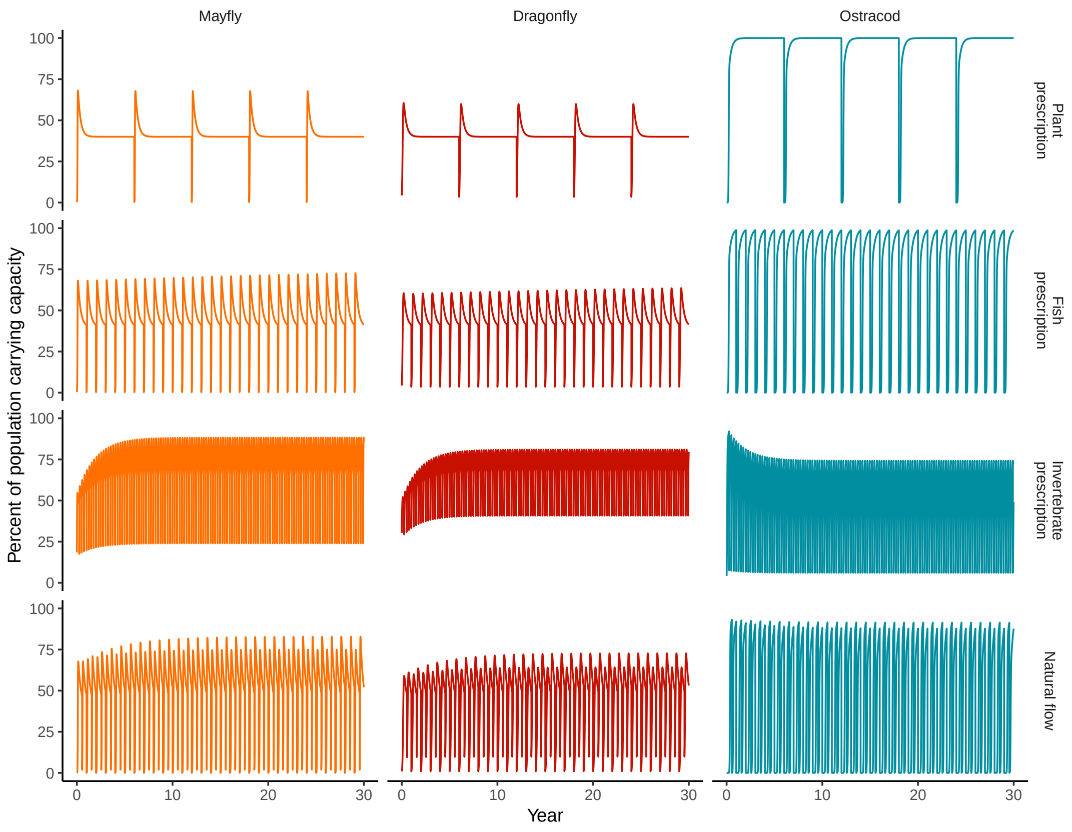


**WebFigure 3:** Results of simulations for the three invertebrate taxa and four flow scenarios. Maximum mean percent of population carrying capacity for the mayfly and dragonfly was the flow target.

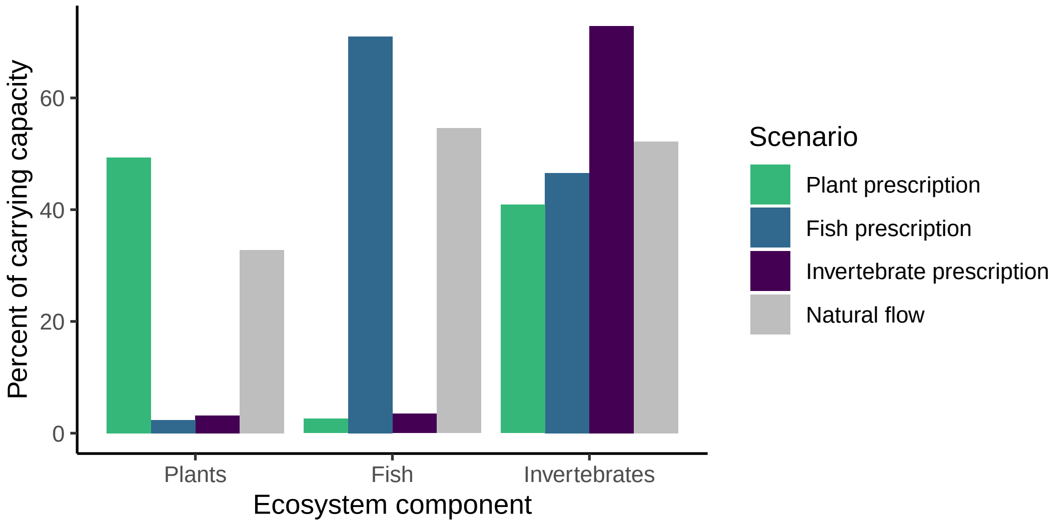


**WebFigure 4:** Results of simulations for target and non-target ecosystem components. These values were used to create the chord diagram in the main text.

### JD Tonkin *et al.* – Supporting Information

**WebTable 1:** Summary of the models used, the final designer flow for each ecosystem component, and the magnitude of the high-flow pulses.

| Group | Model | Flow | Magnitude ($m^3 s^{-1}$) |
|---|---|---|---:|
| Vegetation | Coupled matrix pop. model | 6-y return on large spring floods, preceded by 1 drought and 4 non-event years | 19.8 |
| Fish | Coupled matrix pop. model | 1 large spring flood every year | 19.8 |
| Invertebrates | Time-varying logistic model | 4 small pulses per year | 4.0 |
